# Beyond the rule of seven: translating time across humans and dogs shows diverse clocks set the pace of development and aging

**DOI:** 10.64898/2026.11.13.766441

**Authors:** Christine J. Charvet, Taylor Turner

**Author notes:** Correspondence: Christine Charvet.

## Abstract

Although a dog’s year is often converted to human age by multiplying it by seven, there is little scientific evidence to justify this calculation. Here, we assembled 11,365 observations to generate age alignments across multiple organ systems from humans and 11 canid species and 171 dog groups. We used a range of metrics, including behavioral milestones (e.g., age of eye opening), age-related changes in blood chemistry profiles, disease occurrence, and reproductive traits to translate ages across species and breeds. Our analyses revealed that the pace of development and aging varies across life stages and systems. For example, we found that dogs progress through postnatal development relatively more rapidly than humans. We also found that many biological processes vary relatively independently of each other across and within canid species. Models incorporating system-specific heterochronies across dog breeds outperformed models that assumed breeds differ only in their overall rates of development and aging. These findings demonstrate that estimating a dog’s human-equivalent age by a factor, or a handful of traits, oversimplifies a complex pattern in rates of development and aging. Instead, the pace of aging is dictated from diverse biological clocks.

## Introduction

A common rule of thumb holds that multiplying a dog’s age by seven provides an estimate of its human-equivalent age. Another widely held assumption is that small breeds age more slowly than large ones (Patronek, et al., 1997). However, these ideas have rarely been evaluated within a framework that systematically compares rates of development and aging across organ systems. Here, we integrated biological, behavioral, and life-history measures from humans across 171 dog groups, and 11 canid species to compare the timing of development and aging across organ systems, life stages, breeds, and species.

Dog breeds vary in body size, morphology, lifespan, and disease susceptibility. We leverage this extreme phenotypic diversity to understand the breadth of variation in the pace of development and aging (Wayne, 1986ab; Kemp et al., 2005; Boyko et al., 2010; Drake 2011). We build on a project called *Translating Time*, which is a long-term project aimed to establish age equivalencies across species (Charvet et al., 2025; Cottam et al., 2025; Januel et al., 2026). We collected metrics that track rates of development and aging from multiple organs because biological systems may not mature and age in synchrony (Charvet et al., 2025; Ding et al., 2025).

We use hundreds of timepoints across organ systems to translate ages across species. Some of these changes are rapid; they occur within a matter of days (e.g., eye opening) whereas other processes vary gradually and can span multiple years (Charvet et al., 2025). We used, for example, age-related changes in phosphorus levels, which exhibit similar age-related trajectories across species (Januel et al., 2026). We collected observations from multiple sources, including health records from the Auburn Veterinary Clinic, past studies, and breeder-reported developmental milestones. We used rates of aging and disease incidence rather than lifespan (Ko and Shin, 2025) because lifespan is not a direct biological metric of aging. We found evidence of heterochronies, which are relative accelerations or decelerations in the pace of some processes. This work shows that the pace of development and aging is governed by diverse clocks.

## Results

### The Translating Time Dataset

We collected age-related changes in biological and behavioral changes to capture corresponding ages across human and canid species (Figs 1-9). We define an “observation” as a single datum point (age of eye opening from one or multiple individuals), whereas a “time point” refers to a specific change in biological or behavioral processes averaged by group (species or breed). Some timepoints are based on abrupt changes (e.g., age of eye opening). When possible, we extracted summary statistics, including the minimum, midrange or mean, and maximum ages from a sample. We assumed that the inclusion of these statistics improves the accuracy of age translations. We also extracted timepoints from gradual changes in biological processes (e.g., body growth cessation). In those cases, we used non-linear models to extract corresponding timepoints across groups (e.g., when a blood chemistry profile or body weight plateau; Chang et al., 2016; Puurunen et al., 2022). We also used age-related changes in disease prevalence to align ages across dog breeds. If a dog’s breed was unknown, mixed, or from multiple breeds, their breed was classified accordingly. Some timepoints are classified by sex (age of sexual reproduction, body growth) whereas others are from either or both sexes. For each group comparison (i.e., breed or species), there can be multiple observations per timepoint. We collected information from animals from different environments. When possible, we classified observations according to whether they were obtained from wild or captive individuals. We did not find that the pace of development and aging in the wild varies from those that are captive (y=0.945x+0.02; df=56, Adj R^2^=0.94; p<0.01). The slope of predicted log-transformed timepoints from captive versus wild individuals is not statistically significant (t=-1.76; p=0.08). The data were intentionally collected across different environments to capture variation representative of the species under study. Sample size and number of dog groups vary by analyses.

**Figure 1.**
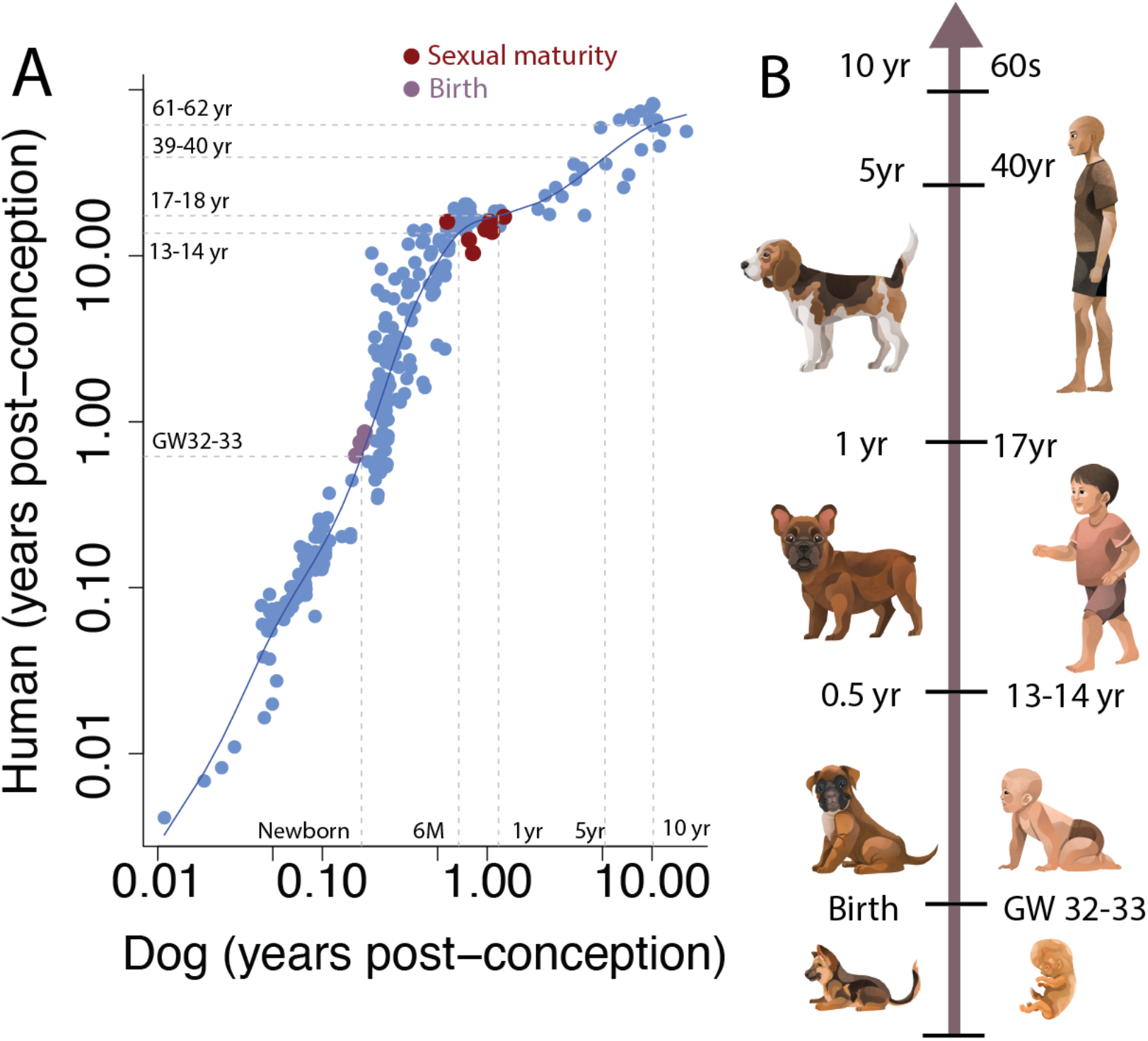
(A) Age matches between dogs and humans are based on corresponding timepoints. (B) According to these analyses, a newborn dog equates to a fetus at gestational week (GW) 32-33, and a 1 year old dog equates to a 17 year old human.

### Human age in dog years

We fit a smooth spline through the log-transformed time points in humans versus dogs to equate ages across species. This model accounts for 95.4% of the variance (n = 298 time points; Fig. 1A). These analyses are based on 5,791 observations and 302 timepoints from 153 different dog groups. On average, there are 20 observations per timepoint in dogs and 5.2 observations per timepoint in humans. We omitted birth (n=4 timepoints) from the smooth spline fit because birth varies independently of other processes, which may impact age translations (Workman et al., 2013). Postnatal development (i.e., the time to reach sexual maturity) is relatively lengthened in humans compared with dogs (Fig. 1A, 2A). Accordingly, a newborn dog corresponds to a 6 month (GW 32-33) old human fetus. A 1-year-old dog corresponds roughly to a human teenager (17-18 years old), and a 10-year-old dog corresponds to a human in their early 60s (Fig 1C). These findings show that specific life stages are stretched or compressed depending on age and species.

Heterochronies are relative changes in the timing of processes (Fig. 1A), and they are especially evident during postnatal development in humans versus dogs. At a given age (e.g., 1 week old dog), the locomotor maturation (e.g., walking) clusters above the smooth spline whereas the onset of sensory function (e.g., eye opening, auditory onset) lie below the spline (Fig. 2A). This means that human babies have opened their eyes and ears to take on sensory information while their locomotor system has yet to mature. In contrast, dog eyes and ears open roughly in synchrony with locomotor behavior maturation. We performed a t test to compare the age of locomotor development versus onset of sensory function to test for significant differences. Whereas the development of locomotion (=85.31 days post-conception: DPC) and sensory function (=81.4 DPC) are not significantly different in dogs (t=1.14, df=20.6, p=0.27), the development of locomotor development (=637.54 DPC) is significantly delayed relative to onset of sensory system functioning (=179.3 DPC) in humans (t=11.46, df = 16.24; p<0.01). Therefore, humans have a relatively protracted maturation of locomotor system development relative to dogs.

**Figure 2.**
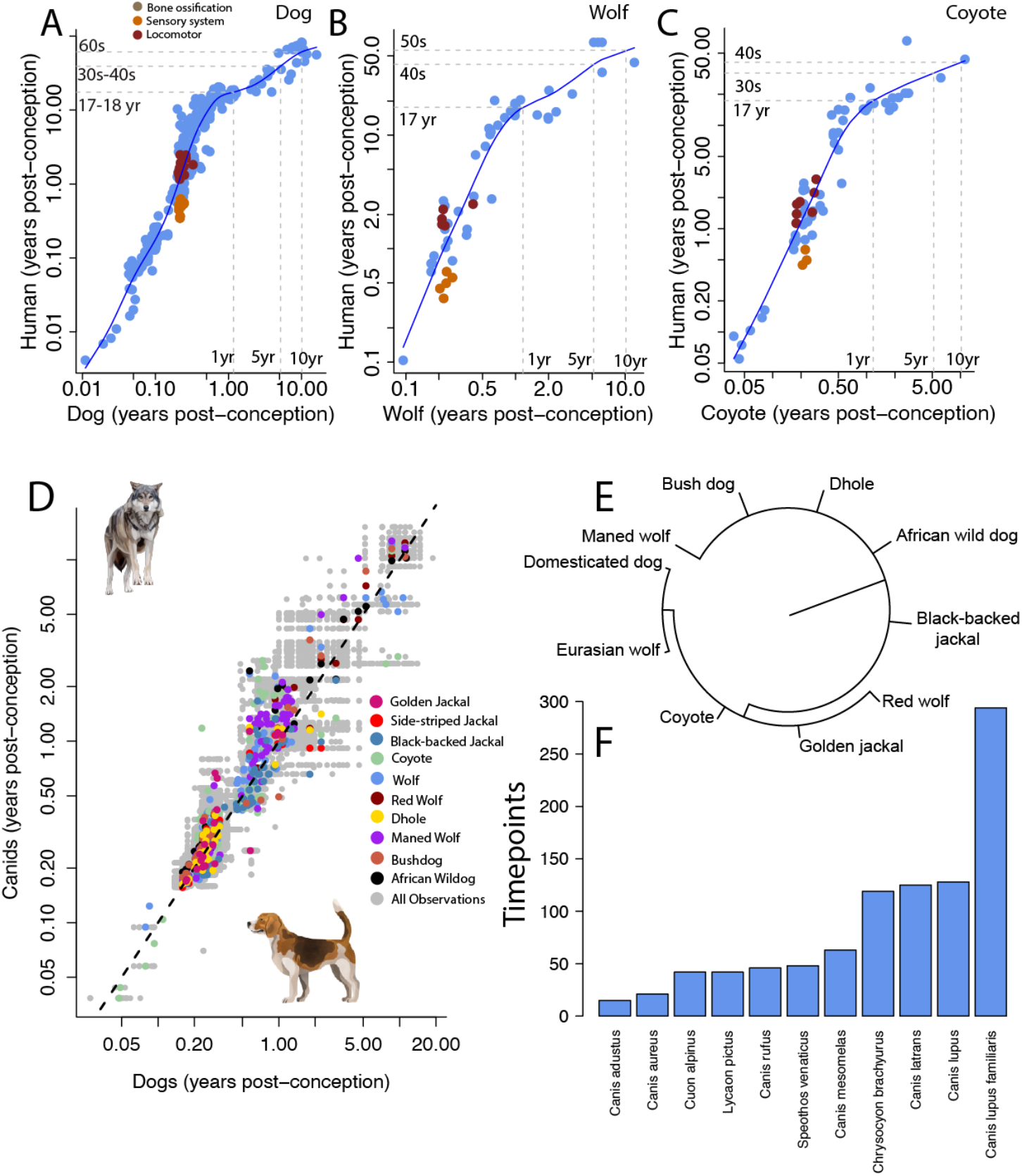
There are relative changes in the pace of development and aging between humans versus (A) dogs, (B) wolves, and (C) coyotes. Smooth splines fit to these data show that human postnatal development is protracted relative to canid species. Also, human locomotor development is protracted relative to onset of sensory system function. In contrast, these processes occur rather simultaneously in (A) dogs, (B) wolves and (C) and coyotes (A-C). (D) Corresponding time points in non-domesticated canid species (n=10) are plotted against those found in dogs. (E) Phylogeny of canid species (F) include wolves and coyotes, which are closest relative to domestic dogs. (F) We have amassed relatively more observations for wolves and coyotes compared with other canid species. Only observations found in at least two species are included in this barplot. (D) The y=x (dashed line) is used to evaluate how the pace of development compares across species. Grey symbols represent all observations across canid species.

### Human age in wolf and coyote years

Human postnatal development is relatively lengthened compared with canid species (e.g., wolves). We fit a smooth spline through log-transformed age of time points in humans versus wolves (Fig. 2B; R^2^=89.7%; n=53 timepoints) and coyotes (Fig. 2C; R^2^=91.9%; n=63 timepoints). We found that humans possess a relatively protracted period of postnatal development compared with canid species. For example, 1 year old wolf equates to an 18-year-old human whereas a 5-year-old wolf corresponds to a 42-year-old human. In wolves as in coyotes, sensory system function and locomotor systems mature roughly synchronously (Fig. 2A-C). Locomotor maturation and onset of sensory system function are not significantly different in wolves (t=0.62; df = 4.55, p=0.56) or in coyotes (t=0.165; df = 7.2; p=0.87). Therefore, the synchronous maturation of sensory and locomotor systems is found across multiple canid species.

### Heterochronies Shape Dog domestication

At first glance, canid species appear to vary more in terms of heterochronies than in their overall pace of development (Figs. 2, 3). In wolves as in coyotes, reproductive traits (e.g., age of sexual maturity, first reproduction) and body growth cessation occur later than in domestic dogs. (Figs. 3, 5) Yet, other processes, bone ossification, occur relatively earlier in wolves compared with domestic dogs (Fig. 3). In the maned wolf, (Fig. 2E), many timepoints occur later than in domestic dogs, suggestive of a protracted period of development (Fig. 3). We first discuss how we developed a pipeline to evaluate how best to characterize variation in timepoints across canid species.

**Figure 3.**
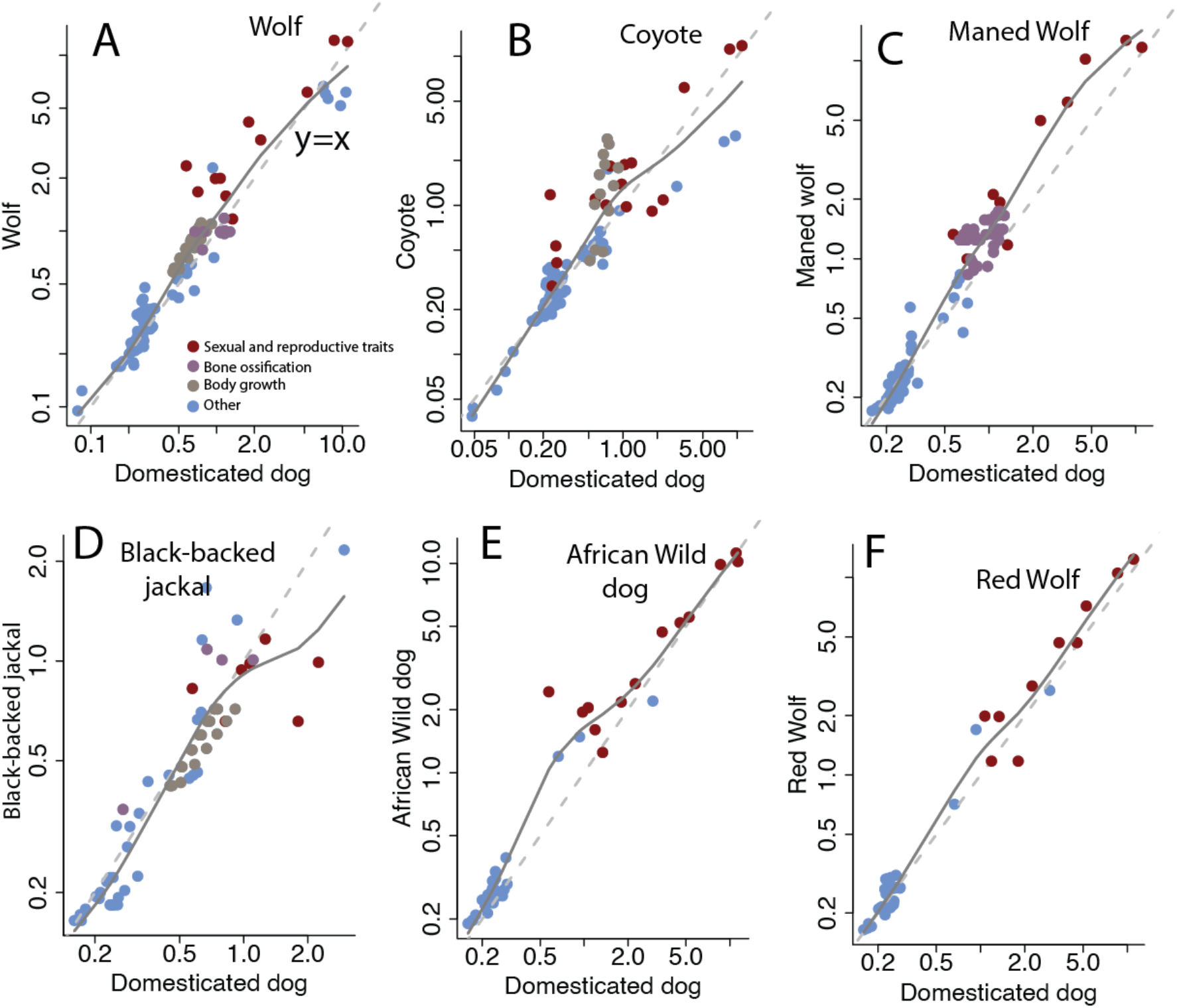
Scatterplots of timepoints across pairs of species. Dashed lines indicate *y = x* and solid lines represent smooth splines fit to the log-transformed data. Time points on the line occur at equivalent ages in both species. Accordingly, some processes, including sexual maturity and body growth, occur later in wolves than in dogs. In the case of the maned wolf, many timepoints lie above the y=x regression.

We built a pipeline to translate ages across species and breeds (Fig 4-9). We collected information across 171 dog groups, but information was not systematically available for every group. To generate an average ordering of timepoints across domestic dogs, we first averaged information across breeds into 10 major dog groups (e.g., retrievers, scent hounds, ancient breeds, and working dogs). Here, included timepoints if they were present in at least 3 dog groups. This filtering step means 789 timepoints with roughly 28-61% of complete data per group were used in the model (Fig. 4D, E-F). We generated imputed datasets with the R package *Amelia*. We then standardized these timepoints by subtracting each timepoint by the earliest timepoint and dividing the resulting values by the difference between the earliest and latest timepoint. This transformation rescaled all observations to a range from 0 to 1, with events occurring early in development having values close to 0 and those occurring later in life receiving values close to 1 (Fig. 4A-B). We used this ordering of timepoints as the predictor variable in the model.

**Figure 4.**
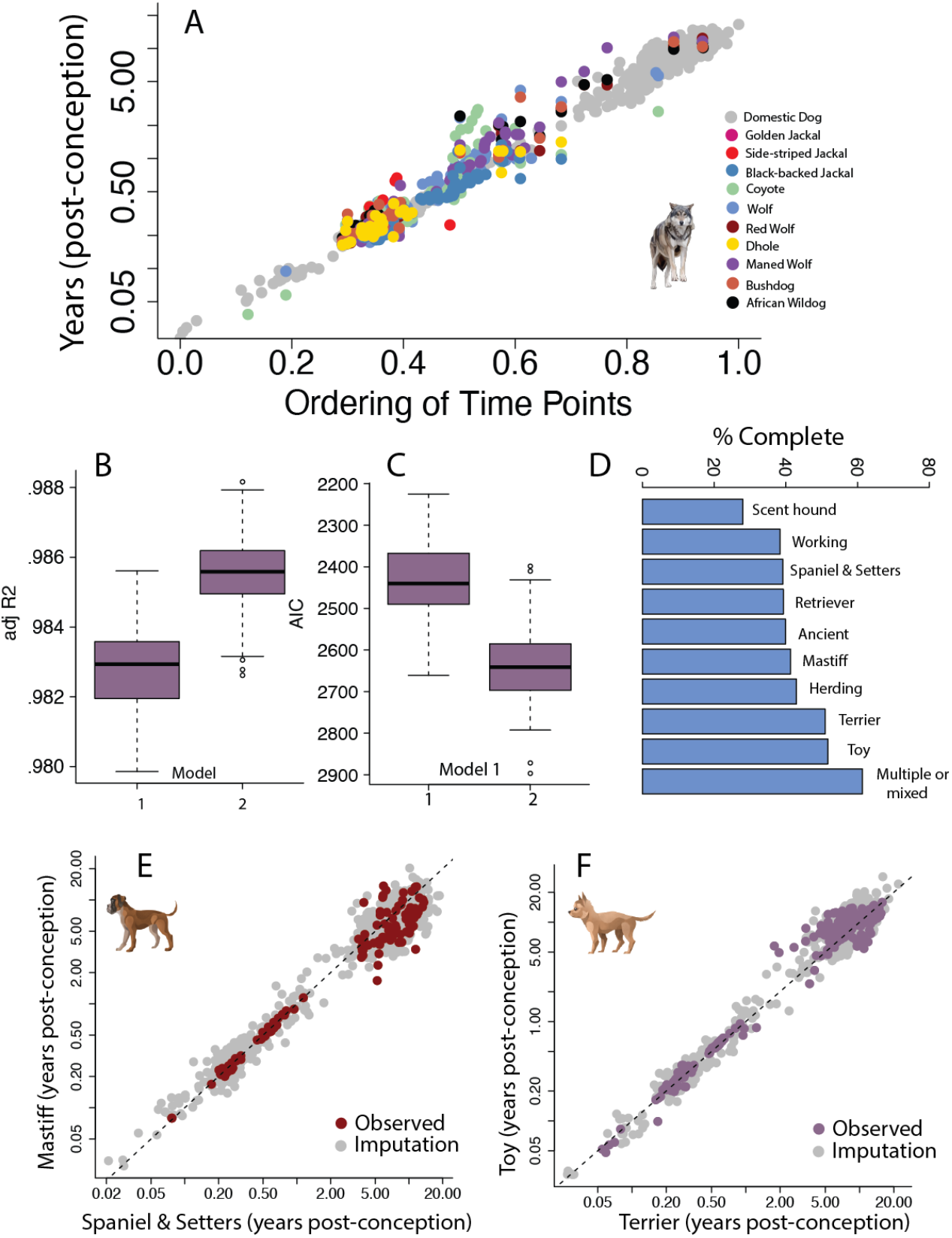
(A) Time points are plotted against a scaled ordering of timepoints. (B) We tested two models to identify which model best accounts for the variance. (B) Both models account for a high percentage of the variance, but the adjusted R^2^ is highest and the (C) AIC score is the lowest for model 2. (D) Percent complete varies by dog grouping. (E-F) Examples of observed versus imputed timepoints for pairs of dog groups. We included y=x as dashed lines. We omitted *Canis aureus* from model testing because there were few timepoints available.

We fit two different models to test whether variation in rates of development (model 1) or heterochronies (model 2) best explain variation across canid species. Body growth, reproduction, and bone ossification were included as factors in model 2. We omitted the Side-striped jackal (*Canis adustus*) in these analyses due to limited sample size (Fig 2F). We tested both models on multiple imputed datasets (n=50) to ensure that findings are robust to variation in the imputed output. The model with the lowest AIC score was considered the best fit model. All models account for a high percentage of the variance (Fig. 4B) but the second model, which considers heterochronies (i.e., bone ossification, body growth, sexual maturity) across canid species accounts for the greatest percentage of the variance and generated the lowest AIC score (Fig. 4C). While wolves and coyotes did not mature significantly more slowly than domestic dogs, there are significant changes in the pace of some processes (Fig. 5). For example, reproduction and body growth are protracted in wolves and coyotes compared with domestic dogs (Fig. 5B-C). Interestingly, the long legs of the maned wolf can be traced back to a protracted duration of bone ossification of its limbs (Fig. 5C). These findings show that canine domestication is characterized by heterochronies in the pace of development.

**Figure 5.**
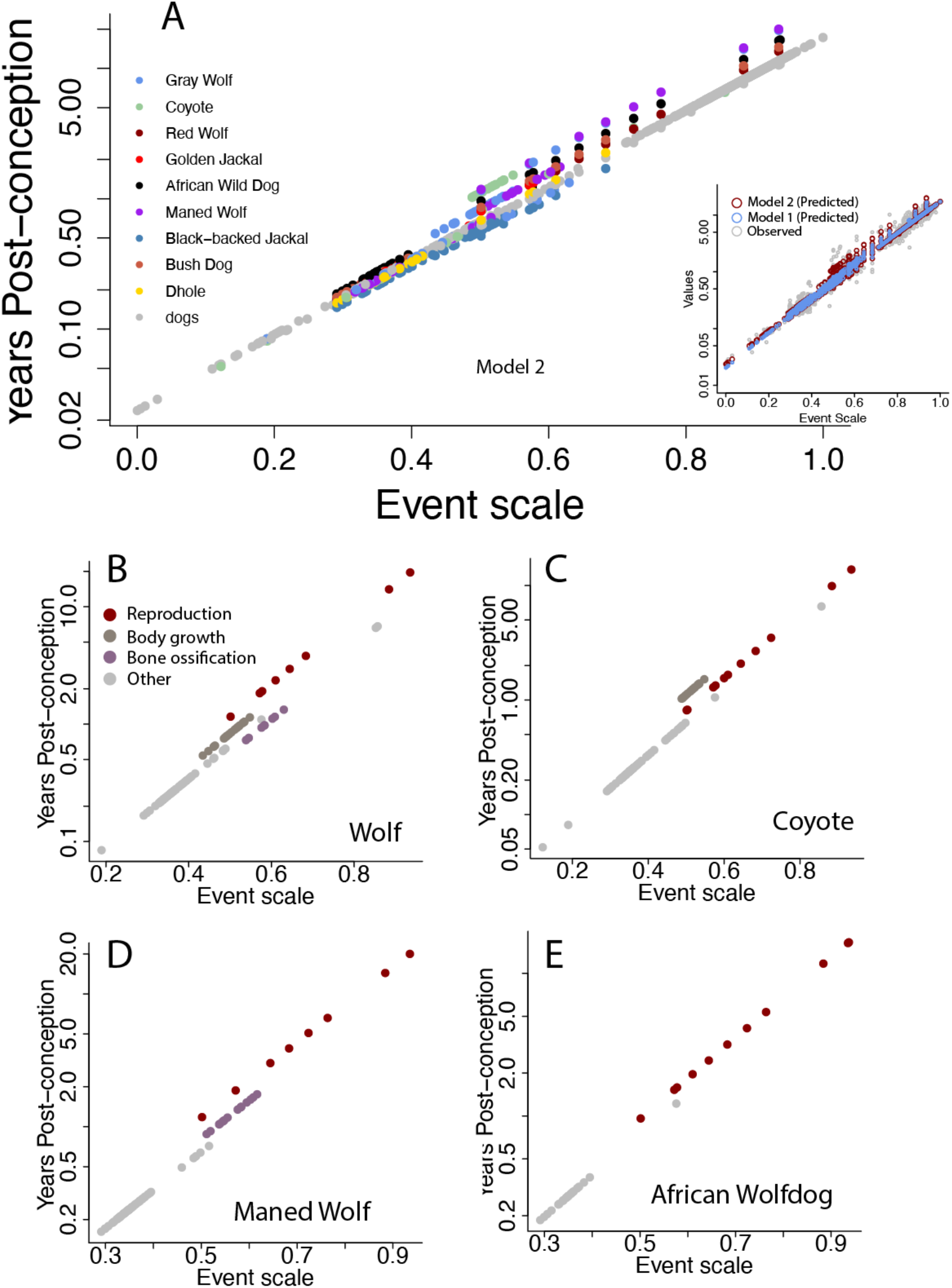
(A) Predicted values are plotted against an average ordering of timepoints generated from dogs. Insert: Predicted values for model 1 and 2 overlap with observed values. Many of the nondomesticated canid species, such as wolves (B), coyotes (C. show a protracted reproductive timeline compared with domestic dogs. (B) In addition, body growth is protracted in the wolf (*Canis lupus*) and the coyote (*Canis latrans*). Bone ossification in the maned wolf (D) is protracted compared with dogs. Sexual maturity is not significantly delayed in the African Wildog (*Lycaton pictus*) compared with domestic dogs.

### Dog breeds age at different rates

We used the same scaled ordering of timepoints as above (Figs. 4-5), and we tested whether timepoints across dog breeds are best characterized by variation in the pace of development and aging (model 1) or by heterochronies (model 2; Figs. 6-8). Because the amount of missing timepoints vary by breed, we iteratively varied the minimum number of breeds (70-125 timepoints per breed) in the model. This effort ensured our results are not swayed by sample size (Fig. 6H, I). We included factors for systems (e.g., cardiovascular, central nervous system; CNS, mass or cancer) to test for heterochronies in model 2. Both models accounted for a high percentage of the variance (Fig. 6I), but the second model resulted in a lower AIC score (Fig. 6H). Collectively, these analyses show that variation in rates of aging across dog breeds are best described as heterochronies.

**Figure 6.**
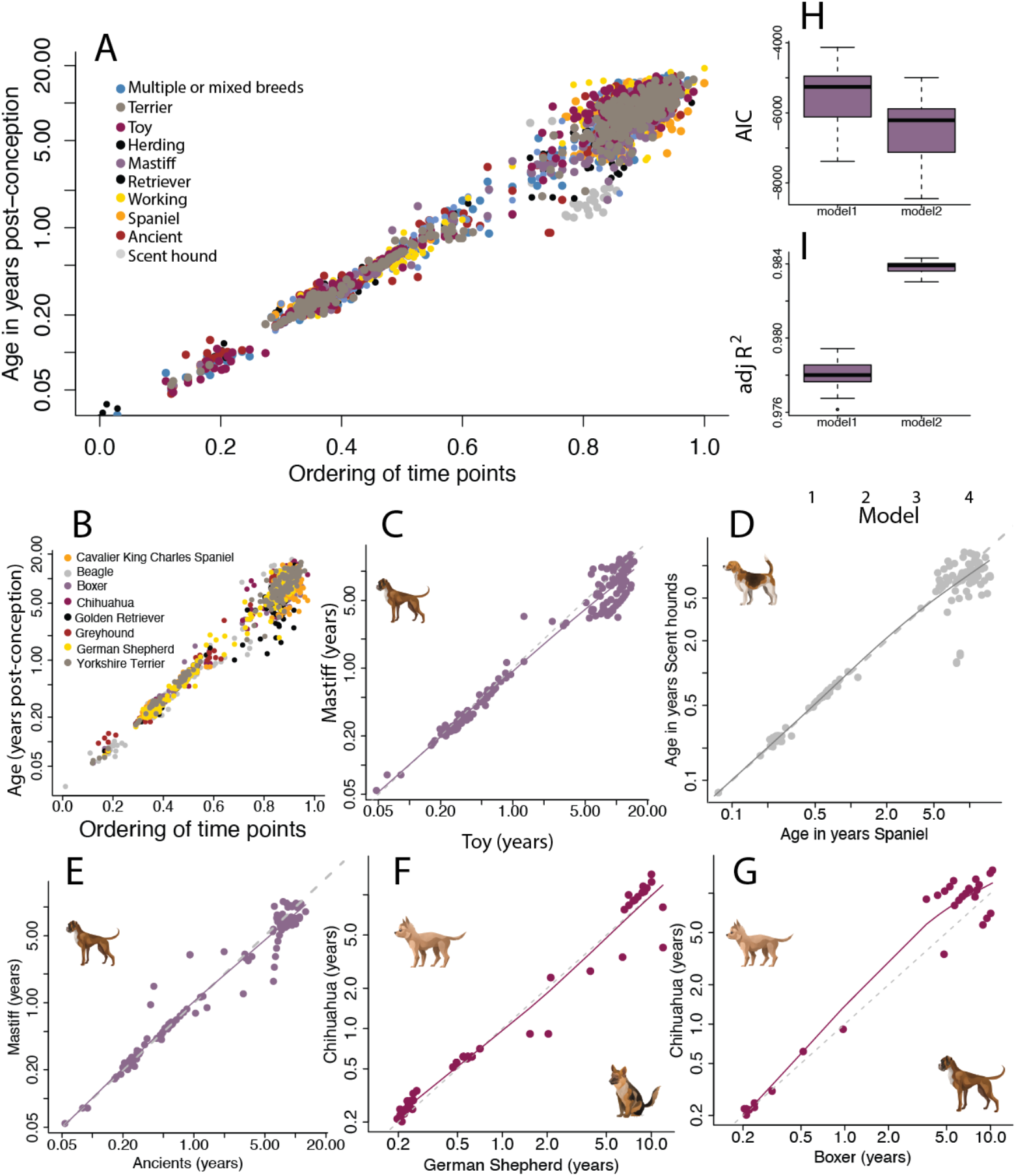
(A) Time points of dog groups (A) and breed (B) are plotted against an average ordering of timepoints. Scatterplots of breed groups (C-E) and breeds (F-G) show overlap in timepoints. Some breeds show variation in rates of aging. For example, many timepoints at late stages of life are protracted in (G) chihuahua compared with boxers. Dashed lines represent y=x. Smooth splines were fit to scatterplots to visualize variation in rates of aging across dog breeds.

While model 2 is a better fit to the data, we consider predicted values from models 1 and 2 as they align with past observations across dog breeds (Figs. 8-9) and because both models account for a high percentage of the variance (Fig. 6I). According to Model 1, toy breeds age more slowly than larger breeds (Figs. 8B-9). According to model 2, breeds differ in their susceptibility to age-related diseases. For example, the accelerated rate of aging in the dachshund central nervous system reflects their early predisposition to intervertebral disc disease and other spinal disorders (Fig. 7A). As another example, boxers show an early predisposition for cancer (Fig 7D, Fig. 9B; Cohen et al., 1974). Some breeds accumulate vulnerabilities across multiple physiological systems (e.g., central nervous system, cardiovascular, Fig. 9) and exhibit an overall accelerated aging trajectory (Fig. 9D). The accumulation of multiple age-related health burdens in some breeds contributes to overall accelerated rates of aging.

**Figure 7.**
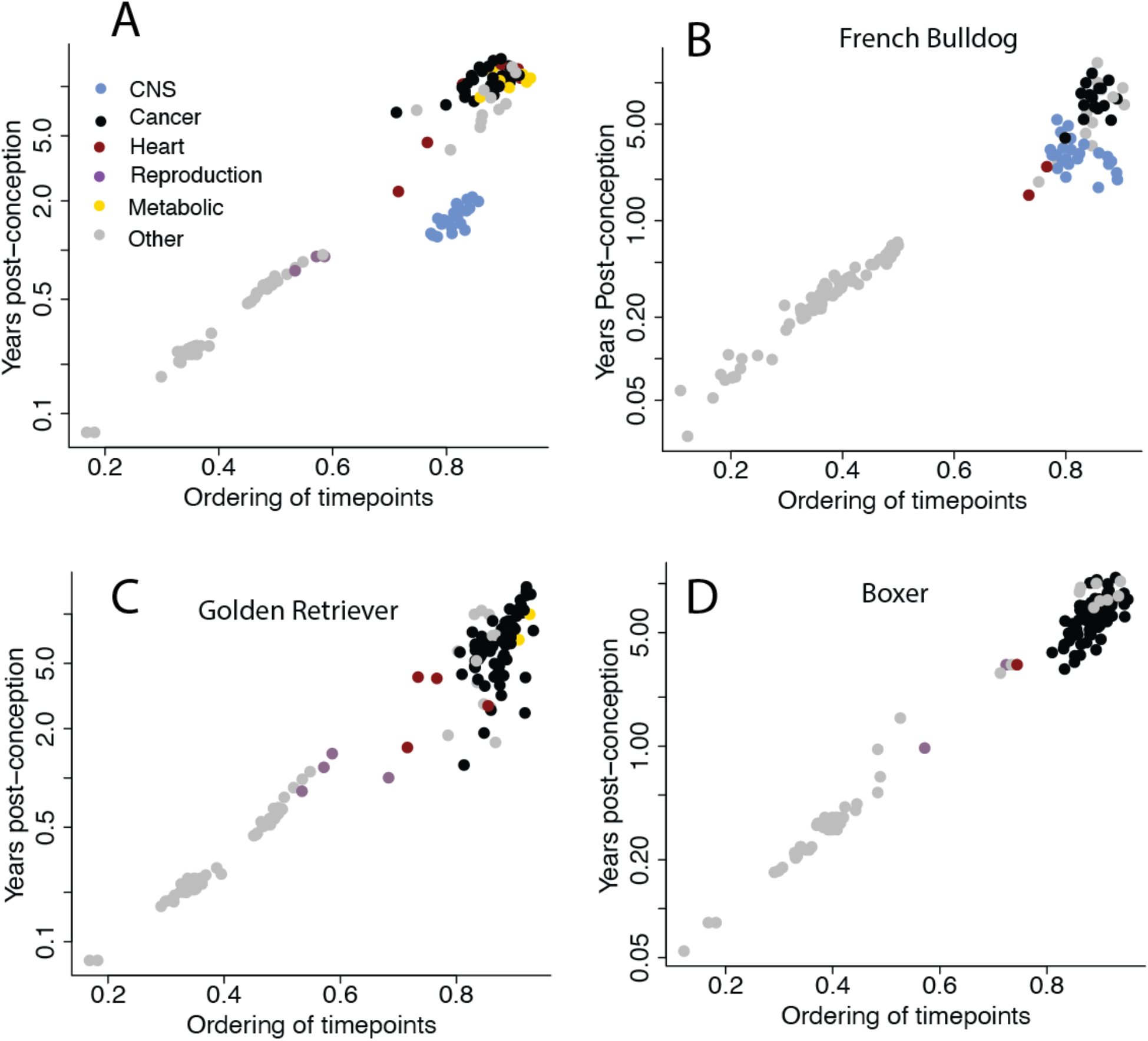
Timepoints in the dachshund (A), the french bulldog (B), golden retriever (C) and Boxer (D) are plotted against the average ordering of timepoints. The dachshund and the French Bulldog have predispositions for early CNS disease. The golden retriever no and boxer show early predisposition for cancer.

**Figure 8.**
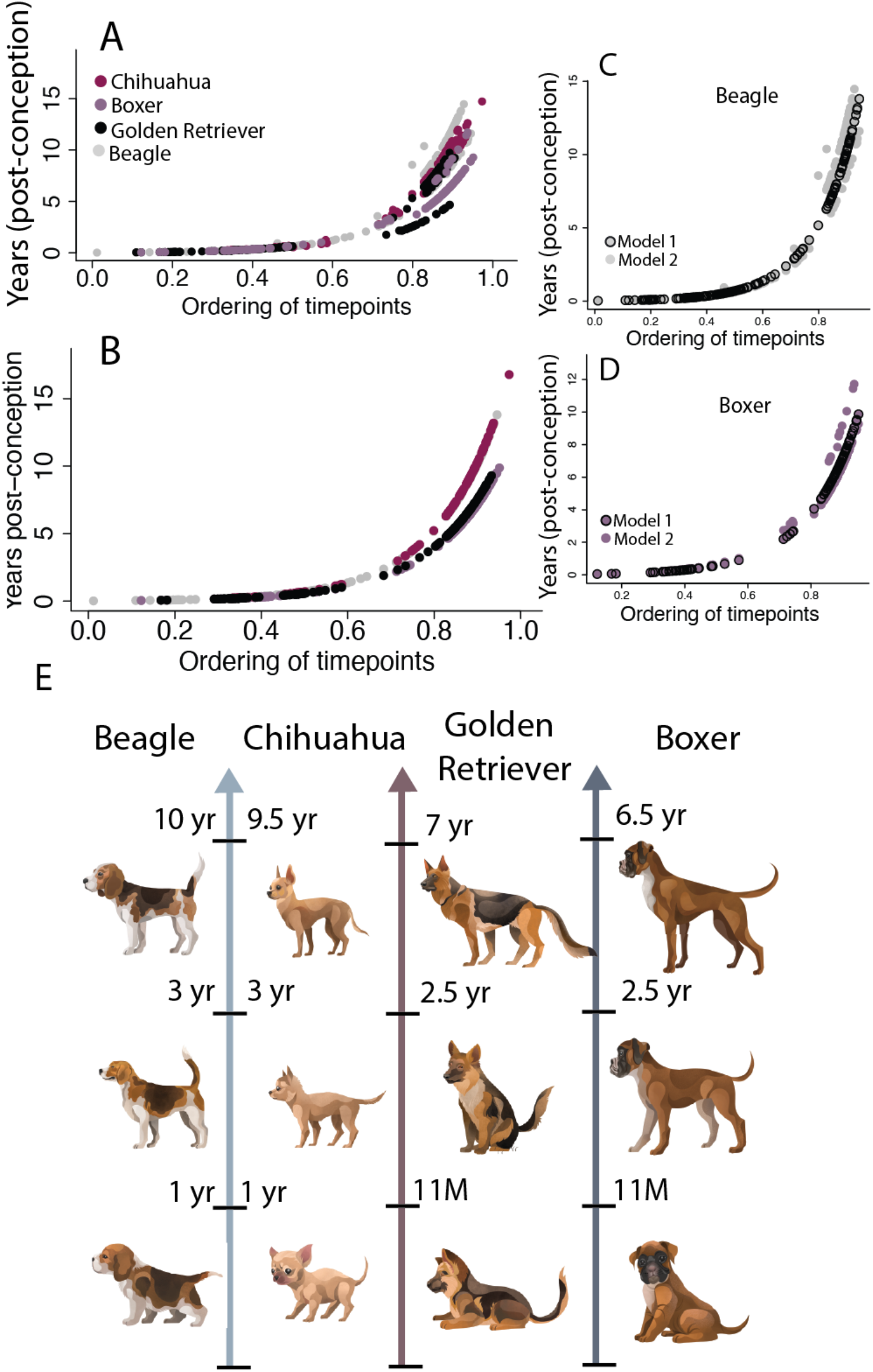
Examples of age alignments across beagles, chihuahua, golden retriever, and boxers based on predicted values from models 1 (A) and 2 (B). Predicted values from both models overlap in the beagle (C) and boxer (D). (E) Approximate age alignments show boxers and golden retrievers age faster than beagles and chihuahua, which were retrieved by plotting data from model 1.

**Figure 9.**
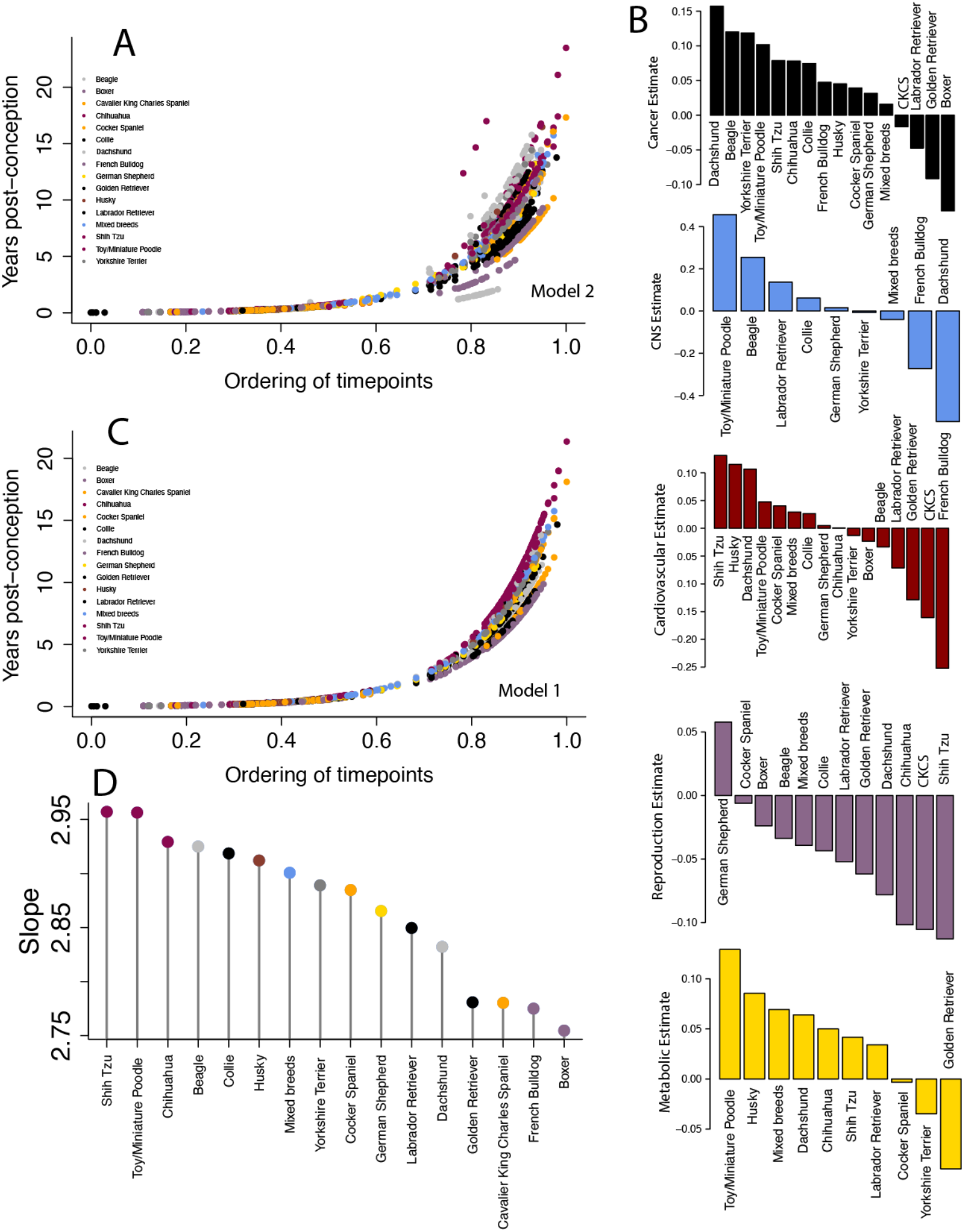
Predicted values from the models 1 (A, B) and 2 (C, D). (B) Heterochronies are factors for breeding by systems. (D) Rates of development and aging from model 1 (D) are color coded according to major dog groups. Here, a minimum of 125 timepoints per breed is used to generate models 1 and 2.

**Figure 10.**
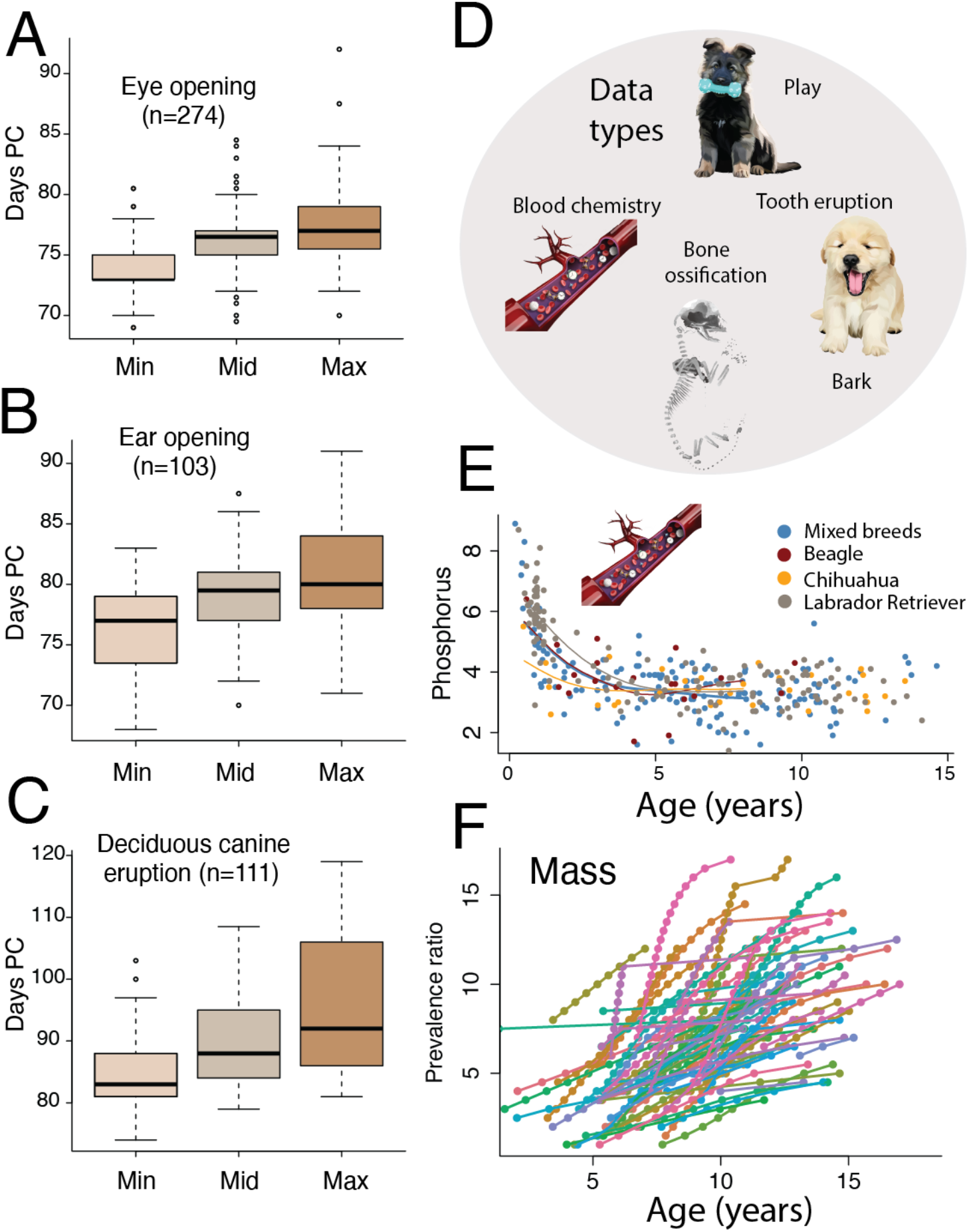
Examples of metrics collected to align ages across groups. For example, we collected data on the age of eye opening (n=274 eye opening observations; A), ear opening (B), and the eruption of deciduous canines (C). In these cases, we collected statistics (e.g., minimum: min, midrange or mean (mid) and maximum, max) when available. (D) We collected a range of transformations from behavior and anatomical changes (e.g., play, bone ossification, bark). In addition, we collected timepoints from gradual changes in biological change (E-F). Phosphorus (E) and age-related changes in prevalence ratios of diagnoses (F) are obtained in the clinic.

## Discussion

We leveraged the extreme phenotypic variation across canid species to generate age alignments across breeds and species. We found that the pace of development and aging varies substantially across species, life stages, and organ systems. We discuss how these results challenge the notion that a human’s age is a multiplicative factor of dogs. We also discuss how our study reshapes our understanding of the impact of domestication on the pace of development and aging.

### Translating time in dogs and humans: moving beyond the rule of seven

The practice of multiplying a dog’s age by seven to estimate a human-equivalent age is no longer tenable. Instead, our results show that canid postnatal development is relatively compressed relative to humans (Figs. 2,3). These observations align with other cross-species comparisons. For example, mice, cats), which possess a relatively compressed postnatal period compared with humans (Cottam et al., 2025; Januel et al., 2026). We also found heterochronies across humans and dogs. For example, locomotor maturation (e.g., crawling, walking) is significantly protracted in humans compared with canid species. These findings are consistent with arguments put forward by Cusack et al. (2024), which argue that a relatively extended duration of locomotor maturation provides a prolonged window to learn the acquisition of complex cognitive and locomotor abilities.

### Variation in rates of development and aging across canid species

We found evidence for changes in the pace of development and aging and for heterochronies across canid species. For example, wolves and coyotes don’t possess a significantly lengthened duration of development, but they do show heterochronies compared with dogs. Some biological processes (e.g., body growth cessation, reproductive maturity) are protracted in wolves and coyotes compared with dogs (Fig. 5). These findings suggest that growth and reproductive timing are evolutionarily labile and contributed to a wide variation in dog size. Collectively, these findings show that heterochronies shaped dog domestication.

### Different dog systems age at different rates

Variation among dog breeds is driven more by heterochronies in biological systems than by uniform differences in rates of development and aging. Some breeds show accelerated rates of aging across many systems (e.g., boxers), while other breeds exhibit relative slowed rates of aging across multiple systems (e.g., shih tzu; Figs. 8-9).

The cumulative balance of these heterochronies shapes breed-specific patterns of aging (Figs. 7-8). Mastiffs (e.g., boxer) are some of the fastest to age whereas toy breeds (e.g., shih tzu) are some of the slowest to age (Figs. 8, 9). These findings are consistent with a large body of evidence showing that many chronic diseases have breed-specific predisposition (Fleming et al., 2011; Donner et al., 2018; Caballero et al., 2026). More work is needed to define the systems that age at different rates.

Past work has relied on lifespan as a proxy for aging in dogs (Patronek et al., 1997), but lifespan is influenced by many factors and may not reflect rates of aging (Cook et al., 2009; Crimmins et al., 2014; Mielke et al., 2014; Austad, 2026). Our past work demonstrated a dissociation between lifespan and rates of aging. For example, translating ages across humans and chimpanzees showed that few chimpanzees live past their 40s or 50s, which equates to humans in their 50s or 60s. These results show that there is a phase in humans past the 70s with no clear counterpart in chimpanzees (Charvet, 2021; Charvet et al., 2023). Life expectancy in humans has lengthened considerably over the past centuries, in part, because of medical care access, which has made the identification of model systems for aging a challenge (Olshansky, 2015; Dattani et all., 2023). Given the dissociation between lifespan and rates of aging, we excluded lifespan from our analyses, and we focused on biological metrics of aging. Our work showed that diverse clocks underlie rates of development and aging.

### Summary

Different systems develop and age at different rates. Approaches that rely on a single metric (e.g., lifespan, blood methylation) overlook substantial heterogeneity across organ systems. By incorporating a broad range of developmental, physiological, and age-related measures, this study translates ages both within and across species.

## Materials and Methods

We collected a total of 11,365 observations across humans and dogs. We subsetted data depending on the analysis performed so that sample size varies by analyses. Some time points, which exhibit sexual dimorphism, are specified by sex (e.g., reproduction, body growth) whereas other timepoints may be drawn from either or both sexes. Some observations are derived from a single individual whereas others are derived from statistics pooled from multiple individuals (e.g., minimum age of eye opening; maximum age of eye opening; Fig. 7). Statistics are included in the definition of each timepoint. For example, the minimum age of eye opening and maximum age of eye opening are distinct timepoints. If information for statistics is not available, we assign the observation as midrange. The inclusion of these statistics was used to improve age translation accuracy.

### Data sources

We used information from the Auburn veterinary clinic, the literature, and online reports from breeders, which report information about the age of developmental milestones (e.g., age of eye opening). We integrate these data with time points from those we previously collected for humans (Workman et al., 2013; Charvet et al., 2022; Cottam et al., 2025; Januel et al., 2026).

### Health records

We used a total of 34,015 visits from 19,700 dogs seen at the Auburn veterinary clinic.

#### Metrics

We used health metrics to generate age alignments across species and breeds. We considered metrics if they vary similarly with age across two or more breeds or species. For example, several variables (e.g., body weight, phosphorus; Fig. 7) increase or decline with age and then plateau (i.e., become relatively invariant; Chang et al., 2016). We used the age of plateaus as corresponding timepoints in the model. To do so, we first filtered outliers (if present) and we fit a smooth spline to the health metrics versus age. We then fit a non-linear model with the library package easynls (model=3) in R. We extracted the age at which these health metrics plateau. We used timepoints if the models reached statistical significance (p<0.05). We fit these models separately to males and females when sample sizes permitted. We extracted observations from age-related changes in blood chemistry profiles from apparently healthy animals that visited the Auburn Veterinary Clinic. We pursued this approach across different populations.

#### Disease Diagnosis

We studied diseases across different systems (e.g., ophthalmic, renal, central nervous system, kidneys) and ages. We limited our analyses to diseases that increase with age (e.g., cataract, arthritis) and that are commonly diagnosed. We considered, for example, cataract and heart failure because age is a factor for such diseases and because we could obtain timepoints from multiple breeds.

Breeds vary in their average lifespan (McMillan et al., 2024). We therefore used prevalence rates rather than age of first diagnosis to control for variation in the age of the population. We calculated the cumulative increases of first disease diagnosis relative to the cumulative percentage of individuals without the specific disease visiting the clinic. We then extracted the ages where animals reach a threshold. For each condition, we separated individuals into disease and non-disease groups. Continuous age data were binned into yearly intervals. For affected individuals, only the age of first disease diagnosis was counted to avoid repeating observations of the same case. Within each age bin, we calculated the cumulative number of newly diagnosed cases relative to the cumulative number of individuals visiting the clinic that did not have the diagnosis. We then fit a smooth spline to model changes in this prevalence ratio versus age. From the fitted curves, we extracted the ages at which specific prevalence ratios were reached. Prevalence thresholds were calculated when there were 5 or more individuals per group, and if there was a continuous increase in disease prevalence with age.

Disease incidence rates can vary widely across populations. Disease prevalence ratios from the Auburn Veterinary Clinic do align with canonical examples of dog breed predisposition (Gelatt and McKay, 2005; Donner et al.., 2018; Puccinelli et al., 2021). For example, we found that dachshunds and the French Bulldog accrue myelopathies at a high rate and that large breeds are at high risk for osteosarcoma. These are well known examples of breed specific diseases. Disease incidence is influenced by many environmental and lifestyle factors. While we report factors for different breeds (Fig. 9B), we caution against over-interpretating these factors because some of these are based on relatively few time points. More work is needed to define these age-related heterochronies.

### Past works

We used past information to extract timepoints. We included age of abrupt transformations (e.g., age of walking, tooth eruption, bone ossification, sexual maturity). We also considered blogs from breeders who reported the age at which animals proceed through milestones (e.g., eye opening, first steps, weaning). We gathered information from humans that were integrated with data we collected previously for humans as part of the Translating Time project (Charvet et al., 2023; Cottam et al., 2025; Januel et al., 2026). Wherever possible, we collected information from individuals across multiple environments (e.g., countries). For example, we compiled multiple traits, such as age at female sexual maturity (n=49), from different continents (e.g., Africa, USA, Europe). We aimed to capture the breadth of variation within human species where possible so that age translations are representative of the human species.

### Age alignments across humans and dogs

The environment varies across species. We collected information from wild and captive canid species. We did not find that individuals from the wild settings aged more slowly than others. Because we have relatively few individuals from the wild compared with those held in captivity, we did not include environment as factors in the model. More work is needed to identify whether environmental factors are associated with differences in rates of aging. Our aim was to encapsulate variation in dogs from their natural environments. We deliberately sampled individuals from different environments (e.g., laboratory, pets) so that our age alignments represent breeds in their natural environments.

### Age alignments across humans and dogs

For cross-species age translations, observations were averaged within species to generate a timepoint for each event. We fit a smooth spline with 10 degrees of freedom to the log-transformed timepoints expressed in years post-conception in humans and dogs. We used a smooth spline rather than a linear model because of the relative compression in life stages across species (Fig. 1).

### Average ordering of timepoints

We developed a model to compare rates of development and aging across dog breeds. To do so, we generated an average ordering of time points that we could use to compare species and breeds. The imputed dataset was generated by first averaging information across dog groups (e.g., retrievers, scent hounds, ancient, working dogs).

We used past work to classify dog breeds (Vonholdt et al., 2010) with the caveat that categorization of some breeds is not fully resolved (Li et al., 2008; Vonholdt et al., 2010; Shannon et al., 2015; Mastrangelo et al., 2018). Designer mixes and breeds that could not be confidently assigned to a specific category were grouped in the multiple and mixed-breed category. This effort aimed to minimize missing information.

### Testing heterochronies

We classified timepoints by organ systems (e.g., cardiovascular disease, n=70; central nervous system, n=45; metabolic disorders, n=24; reproductive, n=10; Fig. 7). These systems were broadly defined. For example, we included blood chemistry data as cardiovascular factors in model 2. Masses and cancers are heterogeneous, but they are included as a separate category. (n=176 timepoints). While our models show that the heterochronies best account for variation in rates of development and aging across canid species (Fig. 6H-I), we caution against over-interpreting breed-specific heterochronies (Fig. 9) because some of these are based on relatively small samples (Fig. 9B). Further work is needed to define the nature and extent of heterochronies across dog breeds.

## Data availability

Datasets to translate ages will be released after acceptance of the manuscript.

## Acknowledgement

This work was funded by Auburn startup funds and Auburn Health Disease Research Fund. The pipeline was not AI generated and every timepoint in the dataset was reviewed by a human. We used AI to edit the flow of sentences and to help flag potential clerical errors while amassing the translating time dataset.

